# Molecular basis for ligand-gating of the human GluD1 receptor

**DOI:** 10.64898/2026.04.06.716743

**Authors:** Anish Kumar Mondal, Haobo Wang, Mae G. Weaver, Iris Zheng, Nikita Kormshchikov, Fairine Ahmed, Edward C. Twomey

**Author notes:** Correspondence (E.C.T.).

## Abstract

The delta-type ionotropic glutamate receptors (iGluRs) GluD1 and GluD2 are ligand-gated ion channels that are fundamental for regulating both excitatory and inhibitory synapses. Rising evidence points to the role of GluD1 in the development of neurological diseases. However, the ultrastructure of human GluD1 (hGluD1) and the molecular basis for its ligand-gating remain unclear. Here, we define the structure of hGluD1 and resolve its ligand-gating mechanism using cryo-electron microscopy (cryoEM) and single channel bilayer recording. While hGluD1 exhibits a non-swapped architecture, it contains conserved iGluR moieties that enable ligand-gating, such as a ligand-binding domain (LBD) tethered to a transmembrane ion channel. Binding of the neurotransmitter γ-aminobutyric acid (GABA) or D-serine to the LBD enables cation influx through the hGluD1 ion channel. Our findings delineate the molecular architecture and function of hGluD1, provide foundations for understanding patient mutations in hGluD1, and will invigorate therapeutic development against hGluD1.

## Introduction

The GluDs are fundamental for organizing and regulating both excitatory and inhibitory synapses^1–4^. While in the iGluR family, GluDs have only recently been defined as ligand-gated ion channels, gating to neurotransmitters such as GABA and D-serine – but not glutamate^5^. However, as with the canonical iGluRs, GluD gating is influenced by and augmented by physiological temperature^5,6^. While trans-synaptic proteins such as cerebellin and neurexin influence the ability of the GluDs to regulate synapses^4,7,8^, rising evidence has pointed to the channel activity of GluDs being regulated through mechanisms related to G-protein coupled receptors (GPCRs)^9–15^.

GluD1 is largely enriched in the forebrain, and influences behavior, memory, and chronic pain through regulating spine development^16–19^. Because of these roles in brain health, dysregulation of GluD1 is linked to psychiatric disorders such as schizophrenia, bipolar disorder, autism, and developmental delay, in addition to intellectual disability and chronic pain^16,20–26^.

The architecture of GluD1 has been characterized as distinct from the other iGluRs, as revealed by cryoEM reconstruction of tetrameric rat GluD1 (rGluD1)^27^. The extracellular domain is comprised of both an amino terminal domain (ATD) and ligand-binding domain (LBD), both of which are arranged into local non-swapping dimer pairs, and a transmembrane domain (TMD) with pseudo four-fold symmetry where the four GluD1 subunits are arranged around a central axis that contains the ion channel.

However, despite this structural information, how disease-associated patient mutations dysregulate hGluD1 and how ligand-gating occurs has been unclear because of the absence of structural and biophysical data on hGluD1. Here, we purified full-length hGluD1, solved its ultrastructure with cryoEM, and characterized its ligand-gating with single channel bilayer recordings. We show how the hGluD1 ultrastructure enables ligand-gating, define how calcium regulates the hGluD1 ultrastructure, how neurotransmitters gate hGluD1, and where patient-linked mutations occur in hGluD1. These findings set foundations for dissecting the regulation of hGluD1, its dysregulation by mutations, and structure-based drug design.

## Results

### Ultrastructure of hGluD1

First, we purified recombinant hGluD1 (Extended Data Fig. 1) and analyzed the hGluD1 ultrastructure in the absence of Ca^2+^ (Extended Data Figs. 2,3). Overall, hGluD1 is quite mobile in the absence of calcium, which is reflected in the overall resolution of the cryo-EM map and local resolutions, ultimately precluding model building. However, hGluD1 maintains an overall “Y” shape like rGluD1. The individual subunit positions in the hGluD1 tetramer (A through D) do not swap across the ATD and LBD, where local dimer pairs are formed between the A-B subunits and C-D subunits. The transmembrane domain (TMD) sits below the LBD.

The individual clamshells of the hGluD1 LBDs are discernible. We describe the clamshell opening of each LBD as the “front” of the domain, and the opposite side the “back.” The C and D subunit LBDs are arranged back-to-back in their local dimer, while the A and B subunit LBDs are arranged front-to-back. Interestingly, calcium was suggested to be critical for the back-to-back dimer arrangement of GluD LBD dimers^28^. While calcium does not appear to alter front-to-back dimer arrangements in full-length GluD2^5,29^, it is possible that calcium locks GluD1 in a conformation where both LBD local dimers are arranged back-to-back^27^.

To test this idea, we prepared cryo-EM samples of hGluD1 in the presence of 1 mM Ca^2+^ (hGluD1_Ca_; Extended Data Figs. 3,4). Based on map quality, calcium limits the overall mobility of the hGluD1 extracellular domain (ECD). The general tetrameric, non-domain swapped arrangement of the receptor is maintained in hGluD1_Ca_, with A-B and C-D subunit local dimers across the ECD. Below the ECD is the TMD, with transmembrane (TM) helices visible within the micelle. While the C and D subunit LBDs are arranged in a back-to-back conformation in local dimer, like the cryo-EM reconstruction of hGluD1 in the absence of calcium, the hGluD1_Ca_ A and B subunit LBDs are now arranged in a similar back-to-back local dimer.

This ultrastructural arrangement is consistent with rGluD1 cryo-EM in the presence of calcium, and with the proposed role of calcium in back-to-back LBD dimer formation in GluDs. Thus, while the hGluD1 ECD samples a conformational space that enables the formation of both back-to-back and front-to-back LBD dimers, calcium may shift this equilibrium to back-to-back LBD dimers.

### Molecular features of hGluD1

There are no full-length models to date of hGluD1, perhaps underscoring the overall flexibility of the hGluD1 ultrastructure. To characterize the molecular features of hGluD1, we performed extensive signal subtraction, symmetry expansion, and local refinements on hGluD1_Ca_ to increase the map quality and enable assembly of tetrameric hGluD1_Ca_ based on the AlphaFold prediction of monomeric hGluD1 (Extended Data Figs. 3-5).While the local cryo-EM maps appear to have over-estimated resolution based on local resolution features at the map edges, the core maps are of adequate quality to enable placement of all molecular features of hGluD1, except for the C-terminal (CT) domain below the TMD and cytosolic linkers between TM helices (Fig. 2a; Extended Data Figs. 3,5).

The ultrastructural arrangement (Fig. 2a) shows that the non-swapped arrangement of dimer pairs is maintained between the ATD and LBD (Fig. 2b,c), as expected from macroscopic features of the hGluD1 cryo-EM maps (Fig. 1) and rGluD1^27^. The map features enable assignment of all TMD features expected from iGluRs, including the pre-M1 helix, M1 TM helix, M2 re-entrant loop helix, M3, and M4 (Fig. 2a; Extended Data Fig. 5).

**Figure 1.**
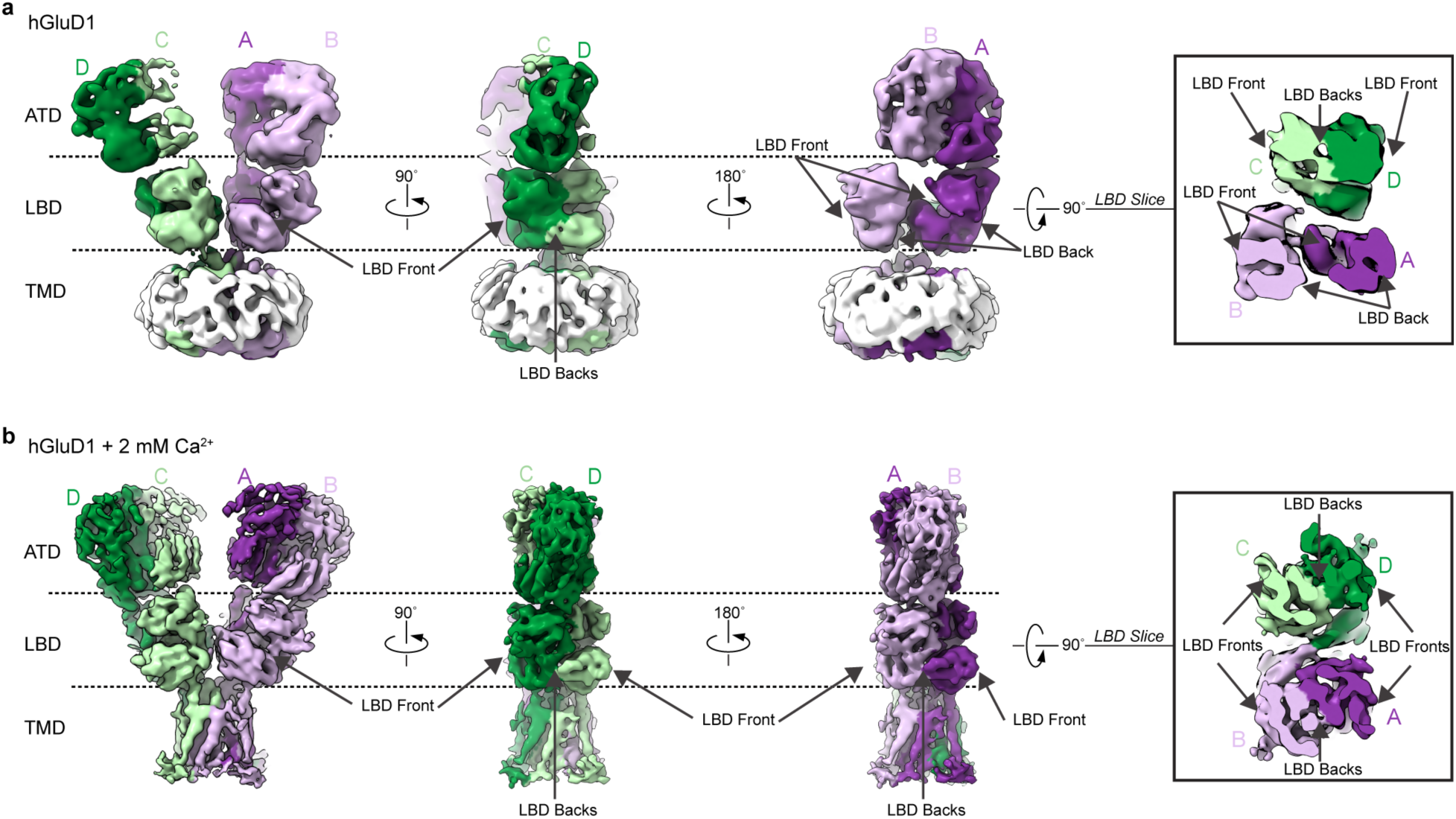
Calcium & the ultrastructure of hGluD1. **a.** Molecular architecture of hGluD1 in the apo state, highlighting the front-to-back and back-to-back arrangements within local LBD dimer pairs. The inset shows a slice through the LBD domains, illustrating the conformational arrangement of the local dimer pairs. **b.** Molecular architecture of hGluD1 in presence of 1 mM Ca^2+^ (hGluD1_Ca_), highlighting back-to-back arrangements within local LBD dimer pairs. The inset shows a slice through the LBD domains, illustrating the conformational arrangement of the local dimer pairs.

The TMD is arranged around a central pore in a pseudo four-fold symmetric manner (Fig. 2d). The top of the pore is lined by M3 helices from each hGluD1 subunit, forming a likely cation channel, as in all other iGluRs, including GluD2. The bottom of the pore is occluded by M2, consistent with its role in selectivity filter formation at the bottom of the iGluR ion channels.

**Figure 2.**
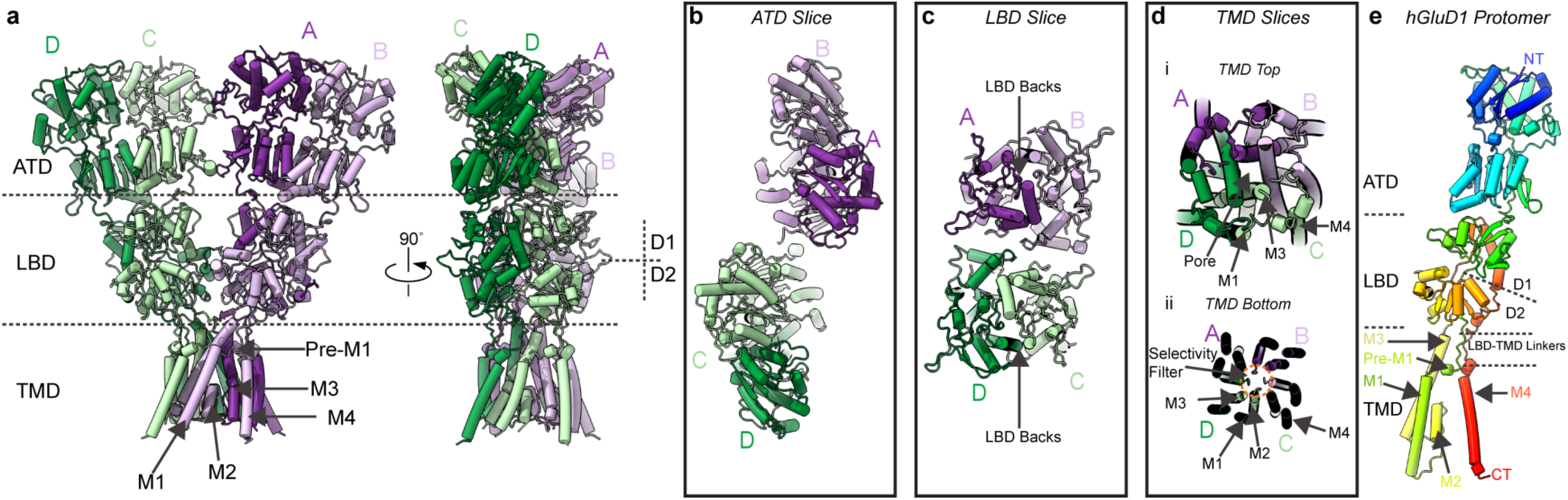
Molecular architecture of hGluD1. **a.** Structure of tetrameric hGluD1_Ca_ **b**. Slice through the ATD domain. **c**. Slice through LBD domains showing back-to-back conformation. **d**. Slices through TMD showing top and bottom views highlighting the pore helices (top, i) and selectivity filter (bottom, ii). **e.** Overview of features in a single hGluD2 protomer from NT (blue) to CT (red).

On the level of an individual protomer, hGluD1 shares complete homology with the other iGluRs: an amino terminus (NT) where the ATD is formed and leads into the LBD. The LBD is structurally arranged into upper half D1 and lower half D2, encoded by two stretches of peptide that form the TMD. Critically, this arrangement enables extensive direct linking of the motions of the LBD (e.g., closure of the LBD clamshell around a ligand) to the TM helices via LBD-TMD linkers (Fig. 2e), a conserved feature in iGluRs for ligand-gating.

### Ligand gating of hGluD1

We reconstituted hGluD1 into brain lipid bilayers to directly test whether we could observe ligand-gated ion channel activity from hGluD1 in isolation. Because physiological temperature (37 <C) augments ligand affinity and open probability in iGluRs^5,6,30^, we performed our recordings initially at 37 <C. GluD1 has been reported to have a role in the regulation of inhibitory synapses through binding GABA^1^. Therefore, we first tested whether GABA elicits canonical ligand-gated cation currents through hGluD1, which based on homology to hGluD2, has a cationic ion channel.

We observe in a concentration-dependent manner that GABA can produce ligand-gated activity directly through hGluD1. In the presence of 1 mM GABA, hGluD1 oscillates between closed (C) and open states (O1 through O4), principally occupying O1 and O2 (Fig. 3a). The current histogram (Fig. 3b) is fit with four curves, representing the subconductance states O1-O4, though the occupancy is primarily O1 (15.2 ± 2.9 pS, standard deviation [S.D.]) and O2 (43.1 ± 4.7 pS). As we step up to 10 mM GABA (Fig. 3c), we observe a qualitative increase in activity and a minor increase in the occupancy of O2 (48.4 ± 5.1 pS; Fig. 3d), consistent with the ligand concentration altering occupancy of O1-O4 in iGluRs^31^.

**Figure 3.**
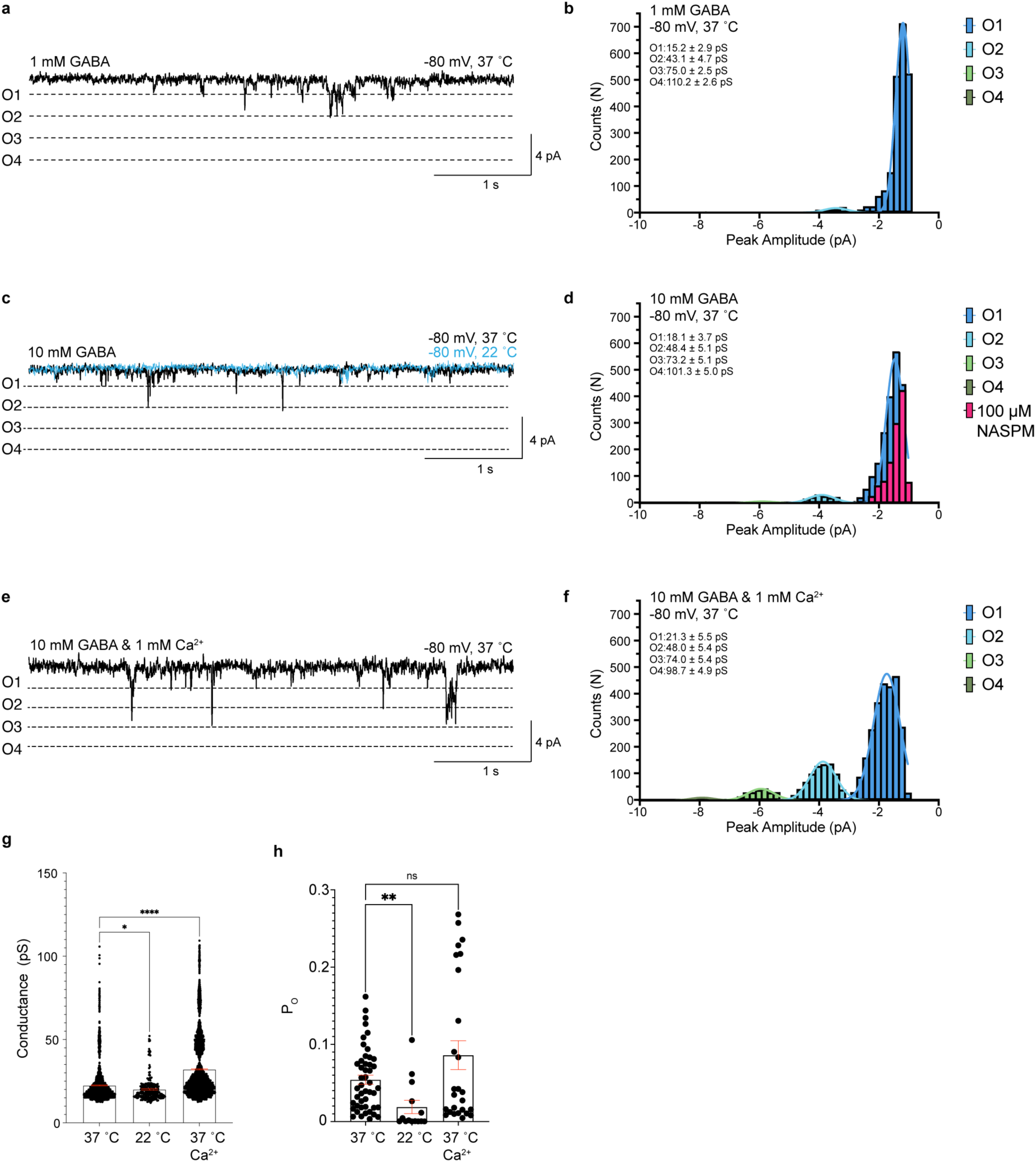
Ligand-gating of hGluD1. **a.** Example of a hGlud1 current trace in presence of 1 mM GABA at 37 ° C. **b**. Current histogram of hGluD1 in response to 1 mM GABA (N = **9** traces), fit with four components at 37 °C (*R^2^* for O1 = 0.99, O2 = 0.88, O3 = 0.84). Subconductance values are represented as mean ± s.d. in the histogram **c.** Example of a hGluD1 current trace in response to 10 mM GABA at 37 °C (N = 12 traces) and 22 °C (N = 13 traces). **d**. Current histogram of hGluD1 in response to 10 mM GABA fit with four components at 37 °C (*R^2^*for O1 = 0.95, O2 = 0.97, O3 = 0.88) and current histogram with co-application of 100 mM NASPM (N = 17 traces). Subconductance values are represented as mean ± s.d. in the histogram. **e**. Example of a hGluD1 current trace in response to 10 mM GABA at 37 °C in presence of 1 mM Ca^2+^. **f.** Current histogram of hGluD1 in response to 10 mM GABA in presence of 1 mM Ca^2+^ (N = 10 traces), fit with four components at 37 °C (*R^2^* for O1 = 0.97, O2 = 0.98, O3 = 0.96, O4 = 0.96). Subconductance values are represented as mean ± s.d. in the histogram. **g**. Overall mean conductance ± s.e.m. of hGluD1 currents at 37 °C, 22.46 ± 0.23 pS (2,285 events); 22 °C, 20.10 ± 0.44 pS (258 events); 37 °C with 1 mM Ca^2+^, 32 ± 0.30 pS (3,639 events). 37 °C versus 22 °C mean conductance *P* value 0.0395, 95% CI of difference 0.09291 to 4.618, d.f. = 6,179; 37 °C versus 37 °C + 1 mM Ca^2+^ mean conductance *P* value <0.0001, 95% CI of difference –10.59 to –8.752, d.f. = 6,179. Columns represent the mean; error bars (red) represent s.e.m. ****P < 0.0001; ns, not significant. **h**. Mean *P_O_* ± s.e.m. of hGluD1 currents at 37 °C, 5.42 ± 0.59 % (47 10s event detections); 22 °C, 1.90 ± 0.86 % (14 10s event detections); 37 °C + 1 mM Ca^2+^, 8.6 ± 1.87 % (26 10s event detections). 37 °C versus 22 °C mean *P_O_ P* value 0.0044, 95% CI of difference 0.01064 to 0.05992, d.f. = 26.53; 37 °C versus 37 °C + 1 mM Ca^2+^ mean *P_O_ P* value **=** 0.2134, 95% CI of difference –0.07783 to 0.01417, d.f. = 30.10. Columns represent the mean; error bars (red) represent s.e.m. ****P < 0.0001; ns, not significant.

Similarly, ligand-gating appears to be temperature sensitive, where channel opening appears to be reduced at 22 <C, consistent with similar experiments on hGluD2^5^ (Fig. 3c). Finally, to verify that the observed currents are from the cationic hGluD1 ion channel, we tested whether the currents are attenuated in the presence of 1-naphthyl acetyl spermine (NASPM), a polyamine channel blocker that blocks GluD currents^5,28,32,33^. Indeed, recordings from hGluD1 in the presence of 10 mM GABA and 100 µM NASPM have reduced currents, reflected by the rightward shift of the current histogram (Fig. 3d), consistent with channel block.

Next, we tested the effect of calcium on hGluD1 channel function because of how calcium alters the hGluD1 ultrastructure. Qualitatively, we observe an increase in the hGluD1 channel activity from 10 mM GABA in the presence of 1 mM Ca^2+^ (Fig. 3e), where there is a general leftward shift in the current histogram (Fig. 3f) with observable occupancy of O3 (74 ± 5.4 pS) and O4 (98.7 ± 4.9 pS).

Overall, we observe a modest decrease in the mean channel conductance from 22.46 ± 0.23 pS at 37 <C to 20.10 ± 0.44 pS at 22 <C (p = 0.0395; Fig. 3g). In the presence of calcium, however, there is a significant increase in the mean channel conductance at 37 <C to 32 ± 0.30 pS (p = <0.0001; Fig. 3g). Juxtaposed to this, the open probability of the channel (P_O_) decreases from 5.42 ± 0.59% at 37 <C to 1.90 ± 0.86% at 22 <C (p =; Fig. 3h). This is reflected by a Q_10_ value of 1.92, consistent with ligand-gating being temperature sensitive in iGluRs. In the presence of calcium, we observe a non-significant increase in P_O_ to <u>8.6 ± 1.87 %</u> at 37 <C (Fig. 3h). This is consistent with observations for hGluD2 in bilayers, and for GluD receptors containing the lurcher mutation^5,28^.

To verify that the observed currents are due to direct GABA binding to hGluD1, we mutagenized the GABA binding pocket. A crystal structure of the isolated hGluD1 LBD shows that R526 is critical for coordinating GABA in the GluD1 LBD^1^ (Fig. 4a), and mutagenesis (R526K) ablates GABA binding. We purified hGluD1(R526K) and reconstituted it into brain lipid bilayers for recording. Indeed, hGluD1(R526K) does not show ligand-gating activity in the presence of 10 mM GABA but can be partially rescued by 100 mM GABA (Fig. 4b). These data show that the ion channel activity we observe is from GABA binding to hGluD1.

**Figure 4.**
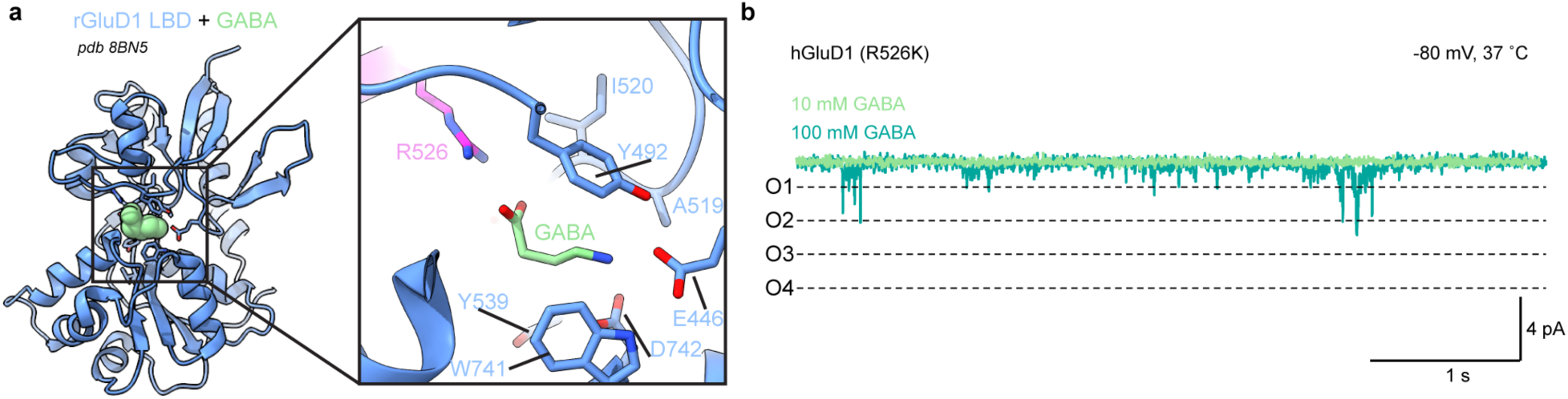
The GABA binding site is required for GABA gating of hGluD1. **a.** crystal structure of isolated GABA-bound hGluD1 LBD (PDB 8BN5), inset highlighting the GABA binding pocket showing R526 residue. **b**. Example of hGluD1 (R526K) current trace in the presence of 10 mM (n = 6) and 100mM GABA (n = 11).

It has also been reported that GluD1 binds the gliotransmitter D-serine^1,32,34,35^. Indeed, both GABA and D-serine elicit ligand-gating from hGluD2. Similarly, we observe that D-serine also gates hGluD1 (Extended Data Fig. 6).

Collectively, this data directly demonstrates that hGluD1 has the intrinsic capability for ligand-gating.

## Discussion

By characterizing the molecular architecture of hGluD1 and through reconstituting purified GluD1 in lipid bilayers for single channel recordings, our data demonstrate that hGluD1, in isolation, functions as a canonical ligand-gated ion channel in the presence of ligands such as GABA and D-serine. Given the definitive data showing that similarly, GluD2 is a ligand-gated ion channel in isolation, a key future direction for the field is how these channels are tuned *in situ* to influence the regulation of excitatory and inhibitory synapses throughout the brain.

Beyond the overall architecture of hGluD1, we observe how cations such as Ca²⁺ regulate the conformational plasticity of hGluD1. Calcium ions are known to stabilize ligand-binding domain (LBD) dimerization and enhance ionotropic activity in the GluD2 Lurcher mutant^28^; however, the structural basis of this calcium-mediated stabilization has remained unclear. We directly demonstrate that calcium induces pronounced conformational changes at the LBD dimer interface by organizing it. Our lipid bilayer recordings support that this arrangement augments channel activity, consistent with previous findings. The mechanisms by which calcium alters GluD2 activity remain unclear.

Through characterizing the hGluD1 architecture, we can map disease-associated variants onto the GluD1 structure (Extended Data Fig. 7)^20^. This reveals that many mutations linked to intellectual disability are localized to the ATD. This observation is consistent with the established role of ATDs in synaptic organization via trans-synaptic partners such as cerebellin-1 (Cbln1) and neurexin-1^36^. However, additional disease-associated mutations are present within the LBD and may also influence agonist binding, highlighting the need for further structural studies of ligand-bound hGluD1. Another region of interest is the A650T mutation in the M3 helix, which is like the A654T Lurcher mutation and may exert similar effects on channel gating. How these mutations alter the hGluD1 ultrastructure, ligand-gating, and *in situ* regulation require future studies.

The interfaces and overall topology of hGluD1 are conserved with rGluD1^27^ (Extended Data Fig. 8). While the overall aligned root mean squared deviation (R.M.S.D) between hGluD1 and rGluD1 is ∼5.4 Å, this is largely due to conformational plasticity due to rotation between the ECD and TMD; each individual domain is nearly identical, with the ATD, LBD, and TMD, having aligned R.M.S.D. values of 1.2 Å, 1.3 Å, and 1.2 Å, respectively (Extended Data Fig. 8).

A limitation of this study is the quality of our cryo-EM reconstruction, which limits our structural discussions to molecular detail. Future work on GluD1 will be required to delineate the exact moieties that enable ligand gating and ion channel permeation. However, given higher resolution reconstructions of hGluD2^5^, the ligand-gating behavior that we report here, and the homology between hGluD1 and hGluD2^4^, we expect the features to be conserved.

Together, our findings provide new insights into the ionotropic function of GluD1 and establish a structural framework for understanding its regulation and disease-associated variants.

## Methods

### Construct design

The human GluD1 (Uniprot ID: Q9ULK0, GRID1_HUMAN) full-length DNA sequence was fused to a TGG linker followed by a thrombin cleavage site (LVPRGS), an enhanced GFP (eGFP), an SGLRS linker, a Strep-Tag II (WSHPQFEK), and a stop codon at its C-terminus. The construct was introduced into a pEG BacMam vector for baculovirus-driven protein production in mammalian cells. To generate the R526K mutant, site-directed mutagenesis was performed on the hGluD1 construct by polymerase chain reaction.

### Protein expression and purification

The hGluD1-containing pEG BacMam vector was transformed into DH10Bac bacterial cells (Gibco, cat# 10301012) to obtain bacmids. The hGluD1 bacmid was then used to transfect ExpiSf9 insect cells (Gibco, A35243), cultured at 27 °C. The P1 baculovirus was obtained by harvesting the supernatant of ExpiSf9 cells after 5 days of transfection. The supernatant containing the P1 virus was added in a ratio of 1:10 (vol/vol) to the Expi293 GnTI^-^ cells (Gibco, A39240), cultured at 37 °C with 5% CO_2_ to initiate the protein expression. Sodium butyrate (10 mM) was then added to the culture to boost the protein expression after 16-18 hours of infection and the cells were moved to a 30 °C incubator. The cells were harvested ∼72 hours post-infection by centrifugation at a low speed of ∼5000g and the pellet was stored -80 °C for further use.

For the purification, the pellet was resuspended in cold Tris buffer (20 mM Tris,150 mM NaCl, pH 8.0) containing a cocktail of protease inhibitors (0.8 μM aprotinin, 2 μg ml−1 leupeptin, 2 μM pepstatin A and 1 mM phenylmethylsulfonyl fluoride). The resuspended pellet was then sonicated with a blunt probe sonicator to lyse the cells (Fisher Scientific, 3-4 cycles of 1 s off, 1 s on, ∼7 power). The lysate was then centrifuged at a low speed of ∼2500g for 20 mins at 4 °C to remove the debris. The supernatant from this sample was ultracentrifuged at ∼125000g for 45 mins at 4 °C to pellet down the membranes. The pelleted membranes were resuspended in Tris buffer, mechanically homogenized and solubilized using a solubilization buffer (20 mM Tris, 150 mM NaCl, pH 8.0, 1% n-dodecyl-β-d-maltopyranoside (Anatrace, D310) and 0.2% brain total lipid extract (Avanti Research, 131101) for 2 hours at 4 °C with constant stirring. The solubilized membranes were then ultracentrifuged (∼125000g, 45 mins, 4 °C) to remove the insoluble materials and the soluble supernatant fraction was incubated with the Strep-Tactin XT Flow resin (IBA, 2-5010) at 4 °C, rotating overnight. The following day, the resin was washed with a GDN buffer (20 mM Tris, 150 mM NaCl, pH 8.0, 100 μM GDN (Anatrace, GDN101) and eluted with a GDN buffer containing 50 mM Biotin (ThermoFisher, 29129). The eluted fraction was concentrated using a 100 kDa molecular weight cut-off filter and centrifugal concentrator down to ∼500 μl in volume. The strepTag II and the eGFP protein were proteolytically removed by incubating the protein sample with thrombin in a 1:100 ratio (w/w) for 1 hour at 25 °C. The protein sample was then further subjected to size-exclusion chromatography using a Superose 6 increase 10/300 column (Cytiva) equilibrated with GDN buffer. The peak fractions from this chromatography were pooled, concentrated up to ∼3 mg/ml and used for cryoEM specimen preparation.

### Cryo-EM sample preparation and data collection

C-Flat grids (CF-1.2/1.3-2Au-50, catalogue no.CF213-50-Au, Electron Microscopy Sciences) were coated with 50 nm gold by sputter coater (Leica EM ACE600) and plasma cleaned (Tergeo Plasma Cleaner, Pie Scientific) to generate holey 1.2/1.3 gold grids with gold mesh. These homemade gold grids were glow-discharged for 2 mins in a Pelco Easiglow (25 mA, 120-s glow time and 10-s hold) just before cryo-freezing the specimen. The purified hGluD1 sample was ultracentrifuged to remove insoluble materials. The protein sample was spiked with 1 mM Ca^2+^ (for the Ca^2+^ bound sample) immediately before cryo-freezing and then 3 μl of the protein sample was applied to the freshly glow-discharged grids in a FEI Vitrobot Mark IV (ThermoFisher Scientific) set at a temperature of 4°C and 90% humidity. The grids were then immediately plunged into liquid ethane cooled by liquid nitrogen. For the Apo state, no Ca^2+^ was spiked. Data collection was performed on either a 300-kV Titan Krios G3i microscope or a 200-kV Glacios microscope (Thermo Fisher Scientific) equipped with a Falcon 4i camera.

A total of 23,427 micrographs were collected for the calcium-bound GluD1 sample. All data were collected on a Titan Krios G3i microscope (ThermoFisher Scientific) operating at 300 kV, equipped with a Falcon4i camera and a selectris energy filter set at a 10-eV slit width. Data collection was performed in an automated manner using EPU software (ThermoFisher Scientific). Data were collected with a total dose of 40.00 e^−^/Å^2^ with a dose rate of 10.89 e^−^ per pixel per s and a pixel size of 0.93 Å per pixel.

For the apo hGlud1without calcium condition, 5,555 micrographs were collected on a 200-kV Glacios Microscope (ThermoFisher Scientific) equipped with a Falcon 4i camera. The micrographs were collected with a total dose of 40.00 e^−^/Å^2^ with a dose rate of 9.31 e^−^ per pixel per s and a pixel size of 1.2 Å per pixel. The defocus range was set from -0.8 mm to - 2.5 mm for all the data collection.

### Image processing

CryoSPARC^37^ (v.4.5.3) was used for all aspects of image processing. Final particle picking was performed with TOPAZ. Details can be found in Extended Data Figs. 2-5 and Extended Data Table 1.

### Model Building

An initial predicted monomeric model of hGluD1 from the AlphaFold database^38^ (model AF-Q9ULK0-F1) was used as the starting point for model building. From this, model building, refinement, and structural analysis were carried out using a combination of ChimeraX^39^, ISOLDE^40^, Coot^41^, and PHENIX^42^, as compiled and distributed by the SBGrid Consortium^43,44^.

### Bilayer recording and analysis

The electrophysiological measurement of purified GluD1 was done on an Orbit mini plate form with a temperature control unit (Nanion Technologies), using MECA recording chips (4x 100 µm cavities, Nanion Technologies, 132002). The recording protocol was adapted from previously established methods^5,45,46^. First, the Orbit mini was calibrated with a standard test cell chip (Nanion Technologies) to adjust the base line after hardware/software starts up. A MECA chip filled with recording solution (HEPEs, pH 7.5, 150 mM KCl) in its holding chamber was then plugged into Orbit mini. A thin lipid bilayer was painted over the recording cavities located at the bottom of the chamber, by dispensing lipid-covered air bubbles on top of each cavity from a 10 µl tip previously dipped in the lipid stock. The lipid stock was 100 mg/mL brain extract total (Avanti, 131101C) dissolved in decane (Sigma-Aldrich, D901-100ML). The over capacitances of the painted bilayer were kept between 10-25 pF. The purified apo WT GluD1 or the R536K mutant receptor was diluted 1:10,000 times from the stock (∼ 4 mg/mL) with the recording solution. A total of 4 µL of diluted protein was added directly on top of each painted cavities using a 10 µL pipette. A varying amount of ligand (GABA or D-serine) was added to the solution before the recording was initiated. The recording solution was also supplemented with 1 mM calcium when indicated. For the blocking condition, the 100 µM NASPM was co-applied with ligand. All the recording was conducted with -80 mV holding potential and a maximum applied current of 200 pA at 37 °C or 22 °C. The data was recorded at 20 kHz sampling rate using Elements Data Reader 4 (EDR4, Elements) software v.1.7.1.

The recorded traces were analyzed using Clampfit software version 11.3 (Molecular Devices). These traces were filtered with an 8-pole Bessel filter with a −3 dB cut-off frequency of 100 Hz and adjusted for baseline before analysis. Five detection levels (−2, −4, −6, −8 and −10 pA) were set for the Single-Channel Search function in Clampfit to identify and measure current events, with the 50% amplitude crossing method used for idealizing current peaks^47^. To minimize local baseline drift during event searching, automatic adjustment for baseline and simultaneous update for all levels (with a 10% level contribution) function was used. Any level change lasting less than 2 ms was excluded from the event picking. In addition, a 10-s detection window was utilized for all the event picking to standardize the sequential calculation and statistical analysis. The picked events were further sorted by detection levels in Clampfit before further being processed in Microsoft Excel. The mean and individual conductance values were calculated using the equation (G) = event current peak amplitude (I_peak_)/holding potential (V_hold_). The statistical analysis and comparison were done in GraphPad Prism 11. The frequency distribution analysis was performed on the peak amplitude and the conductance of all the picked events to generate initial histograms. Nonlinear regression analysis with Gaussian distribution was performed to generate fitted curves on histograms, using the following default equation in Prism 11:

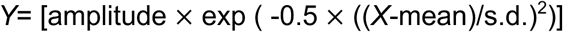

where s.d. is standard deviation. The default rules implemented in Prism 11 to compute the initial values for the Gaussian distribution were defined as the following:

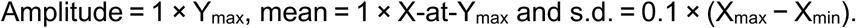

To compare mean conductance and open probability, the unpaired Brown–Forsythe and Welch one-way ANOVA test was used, assuming variances among the experimental groups were unequal. The Game–Howell post hoc test was further used to examine the significance of the difference found among the tested mean values, as the sample size of each group was larger than 50^48^. The overall open probability P(o) was calculated using Clampfit with a default interval setting (100 ms) for each 10-s event detection. Overall, mean open probability was then calculated for each sample condition. The Robust regression and Outlier (ROUT) function in Prism 11 with a default setting of coefficient Q = 1% was used to identify the outliers in each data set^49^.The remaining clean values were compared using the unpaired Brown–Forsythe and Welch one-way ANOVA test in Prism 11. The Q10 value related to P(o) was calculated with the following equation:

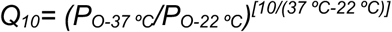

All the final graphs were plotted using Graphpad Prism 11.

### Fluorescence-detection size-exclusion chromatography

The pooled fractions of purified protein samples were further analyzed by fluorescence-detection size-exclusion chromatography^50^ using a high-performance liquid chromatography system equipped with a multi-wavelength fluorescence detector, an autosampler (Shimadzu, SIL-40C), and a Superose 6 Increase 10/300 GL SEC column (Cytiva). The tryptophan fluorescence was used to monitor the protein stability and retention time (excitation at 280 nm and emission at 325 nm).

## Acknowledgements

We thank the members of the Twomey lab for helpful discussions during the development of this work. This work was supported by the National Institutes of Health (grant #R35GM154904) and the One Mind Bristol Myers Squibb Rising Star Award via E.C.T. All cryo-EM data was collected at the Beckman Center for Cryo-EM at Johns Hopkins, which is supported by the Arnold and Beckman Foundation, the Howard Hughes Medical Institute, the Johns Hopkins University School of Medicine, and private, anonymous donors. All computational work was supported through the Johns Hopkins Research Information Technology DISCOVERY high-performance computing cluster.

## Author Contributions

E.C.T. conceptualized and supervised the project. A.K.M. and E.C.T. designed the experiments. A.K.M. performed protein expression, purification, cryo-EM collection and processing. A.K.M. and E.C.T. built the models. A.K.M., H.W., M.G.W., I.Z., and F.A. performed the bilayer experiments. H.W., M.G.W., and E.C.T. analyzed the bilayer data. A.K.M. and N.K. designed the mutant hGluD1. A.K.M. and E.C.T. wrote the manuscript, which was edited by all authors.

## Data Availability

The cryo-EM reconstructions are deposited into the Electron Microscopy Data Bank (EMDB). All structural coordinates are deposited in the Protein Data Bank (PDB).Accession codes will be available upon publication. All electrophysiology data is included in the manuscript and in the extended data.

---

**Extended Data Figure 1.**
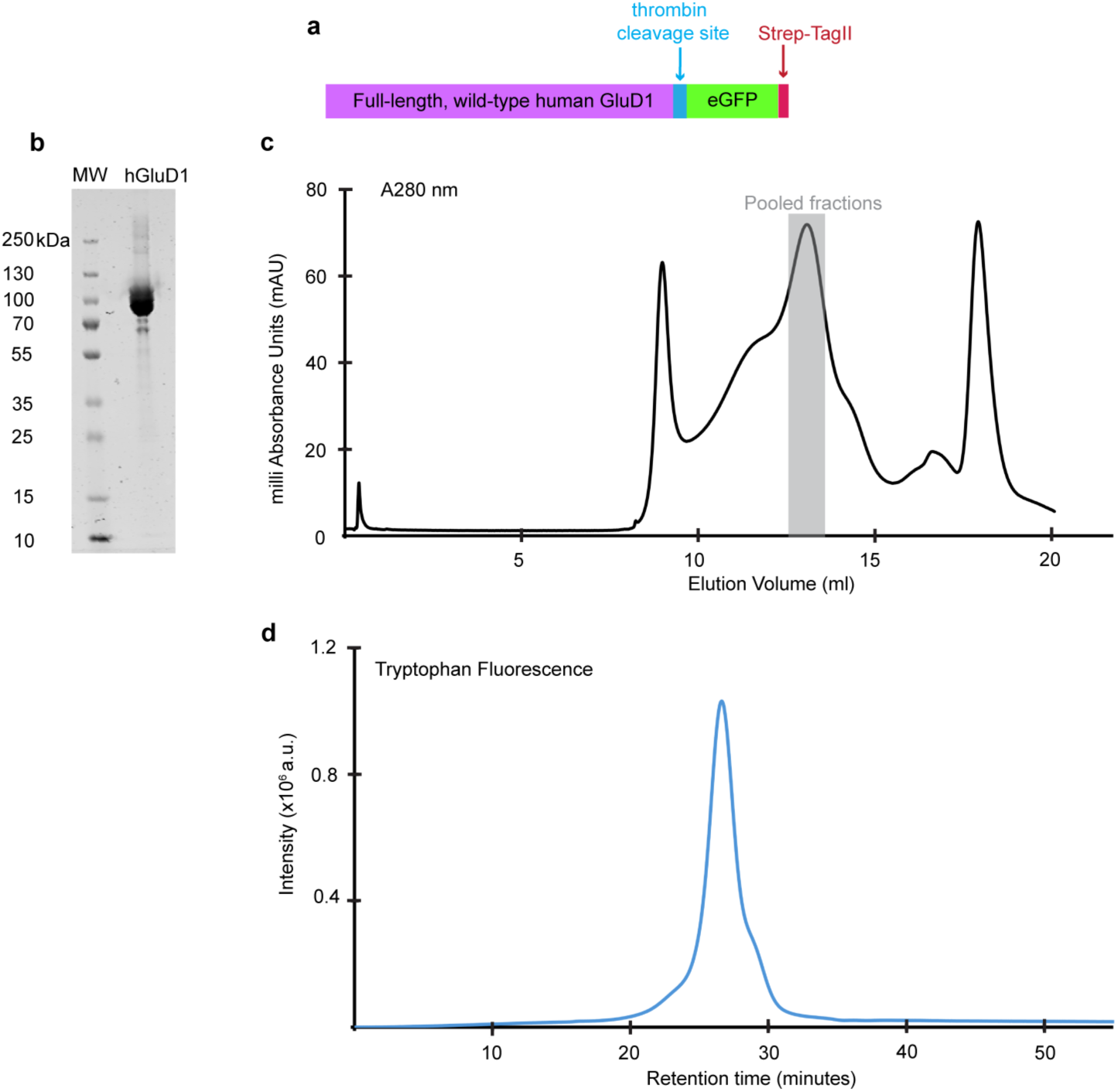
Purification of hGluD1. **a.** Construct design schematic of hGluD1. **b.** Coomassie-stained SDS-PAGE gel of purified hGluD1, molecular weight marker (left), hGluD1 (right). **c.** Chromatogram of size exclusion chromatography (SEC) for the purification of hGluD1, showing the pooled fractions with a grey bar. **d.** FSEC tryptophan fluorescence chromatogram of purified hGluD1.

**Extended Data Figure 2.**
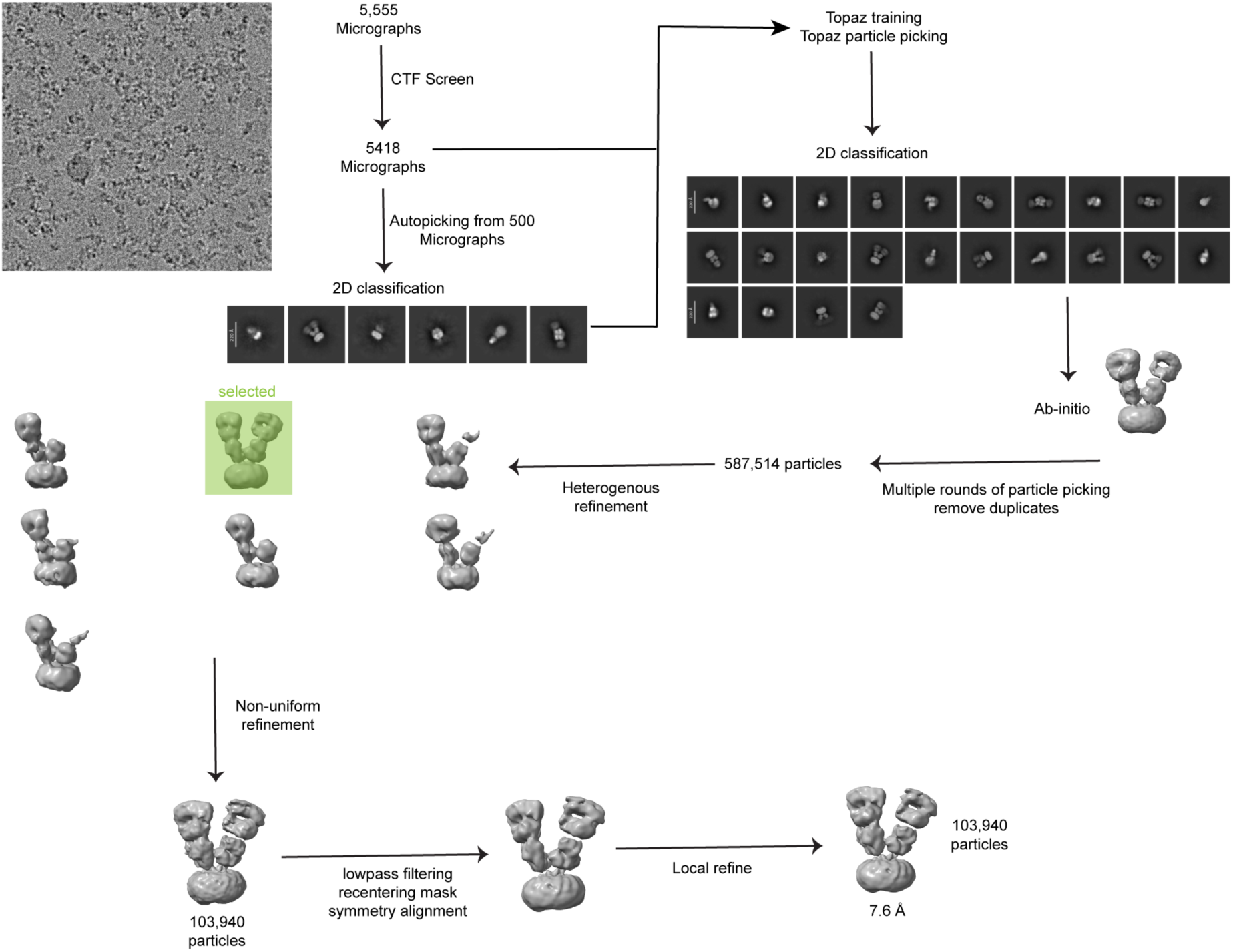
Cryo-EM image processing workflow for hGluD1 (no Ca^2+^) sample.

**Extended Data Figure 3.**
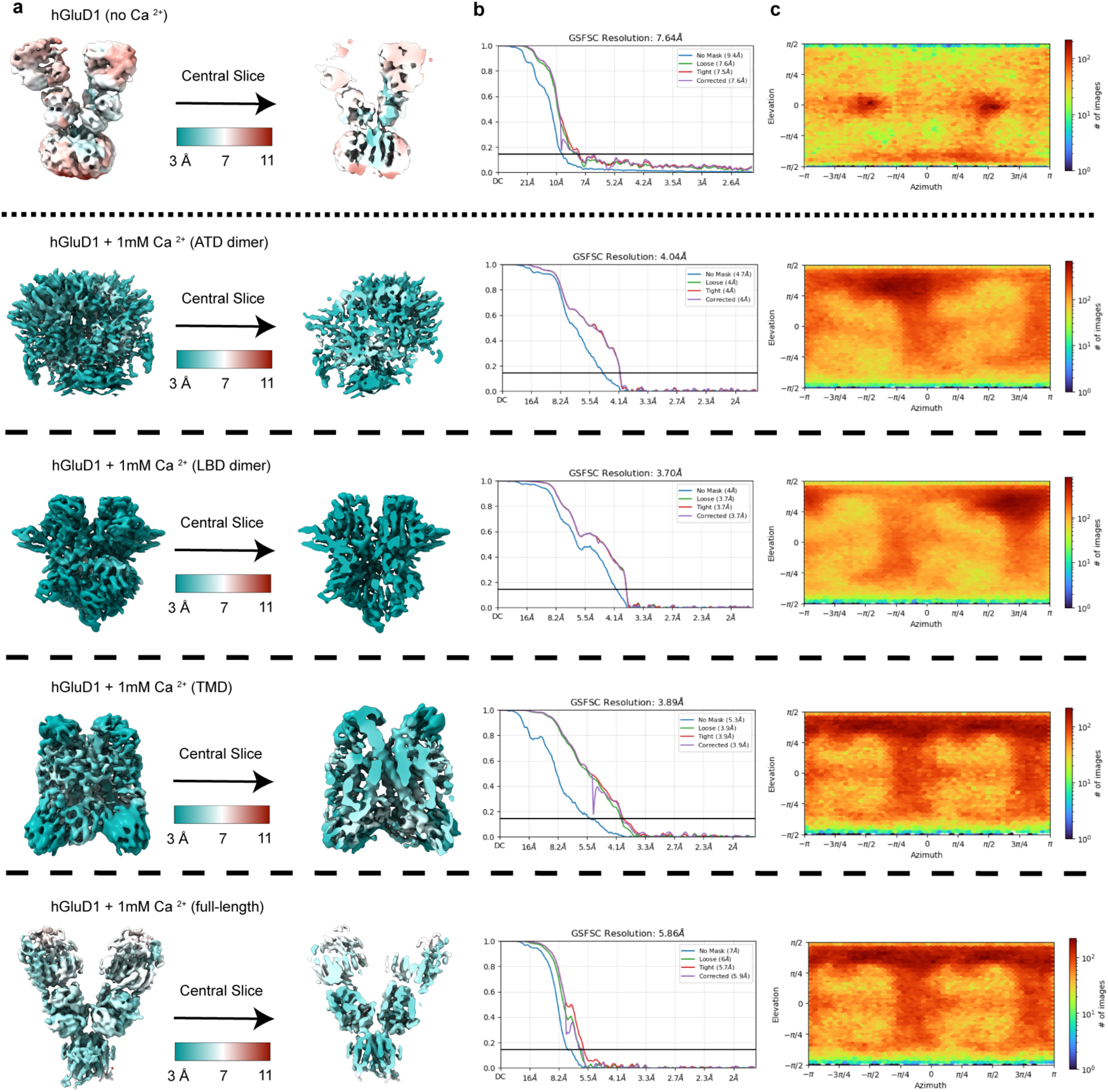
Local and overall map qualities of hGluD1. **a.** Local resolution maps computed for each voxel, computed at Fourier-shell correlation (FSC) = 0.143. **b.** Gold standard Fourier shell correlation (GSFSC) curves for each whole and local map. The black line is Fourier shell correlation (FSC) = 0.143. The Y axis is FSC, the X axis is resolution in Å. **c.** heat map of particle orientation distribution for each map.

**Extended Data Figure 4.**
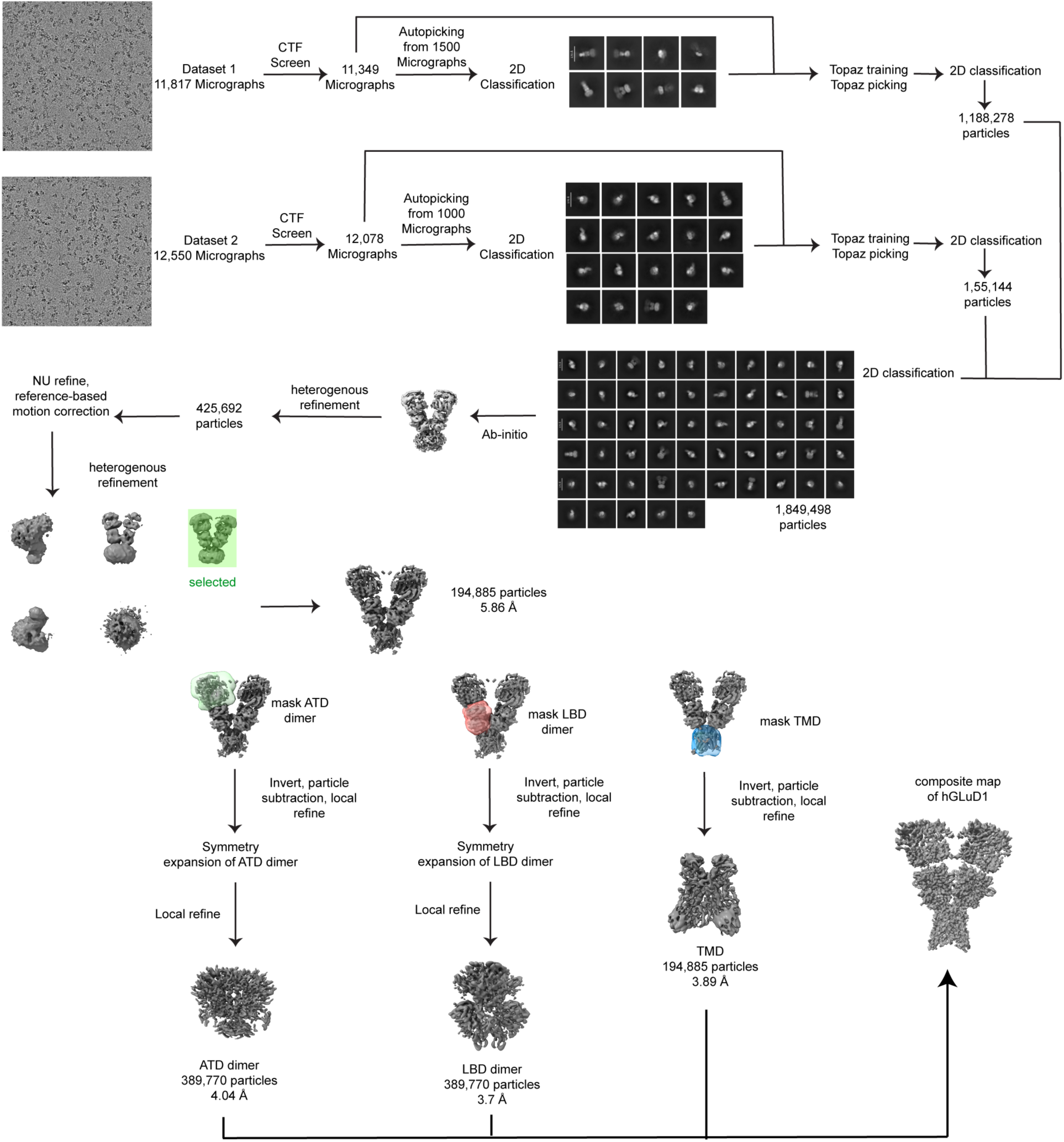
Cryo-EM image processing workflow for hGluD1_Ca_ sample.

**Extended Data Figure 5.**
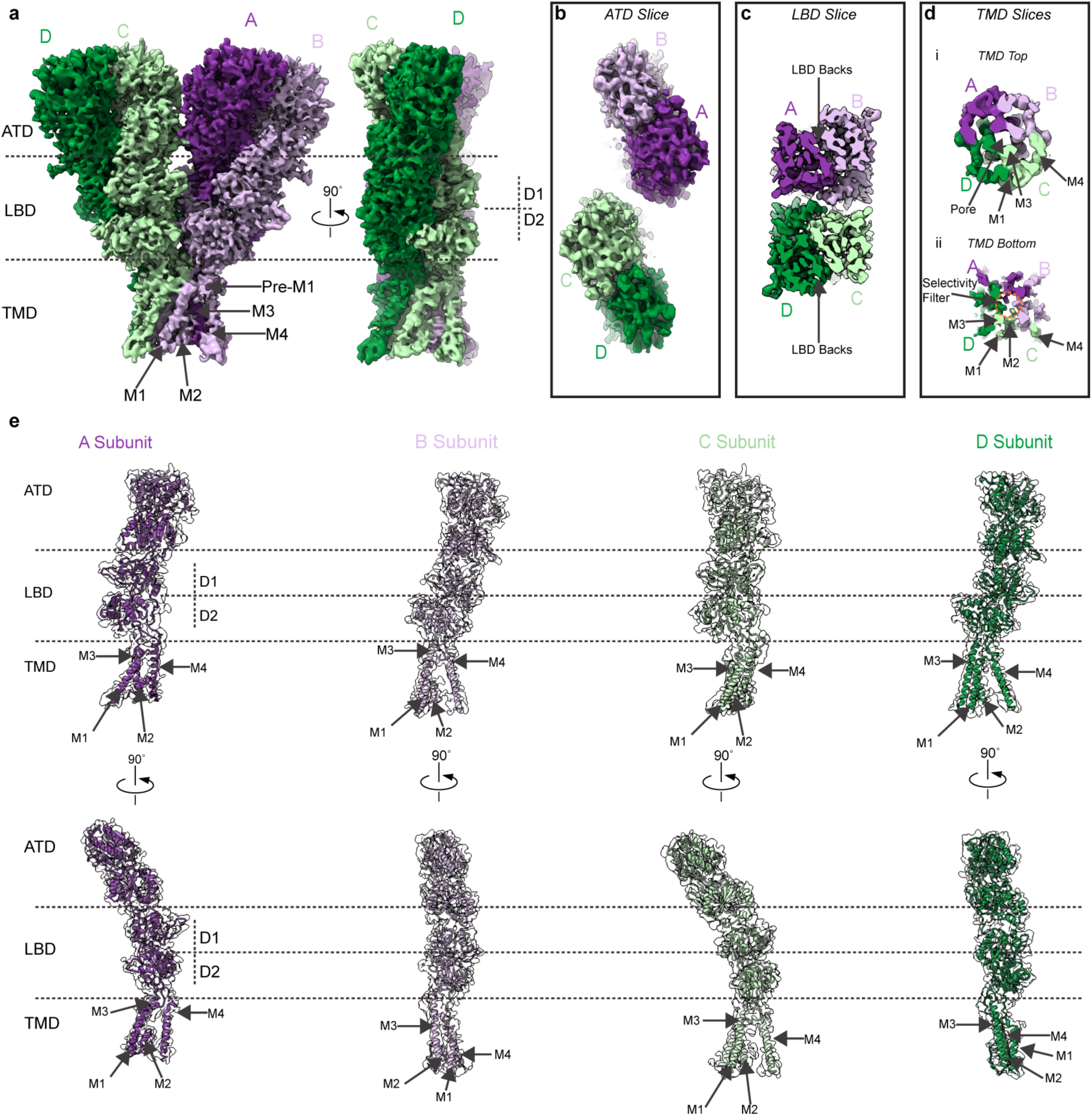
Consensus map, map quality, and model fit for hGluD1_Ca_ data. **a.** Composite map of hGluD1_Ca_ **b**. Slice through the ATD domain. **c**. Slice through LBD domains showing back-to-back conformation. **d**. Slice through TMD showing pore and helices **e.** Subunit protomer fits into the consensus map.

**Extended Data Figure 6.**
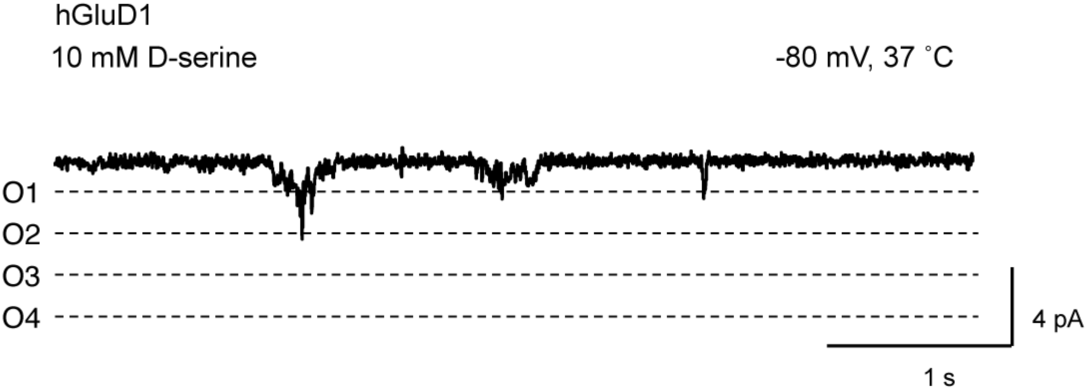
D-serine gating of hGluD1. **a.** Example recording of hGluD1 current traces in bilayers in the presence of 10 mM D-serine (n = 4).

**Extended Data Figure 7.**
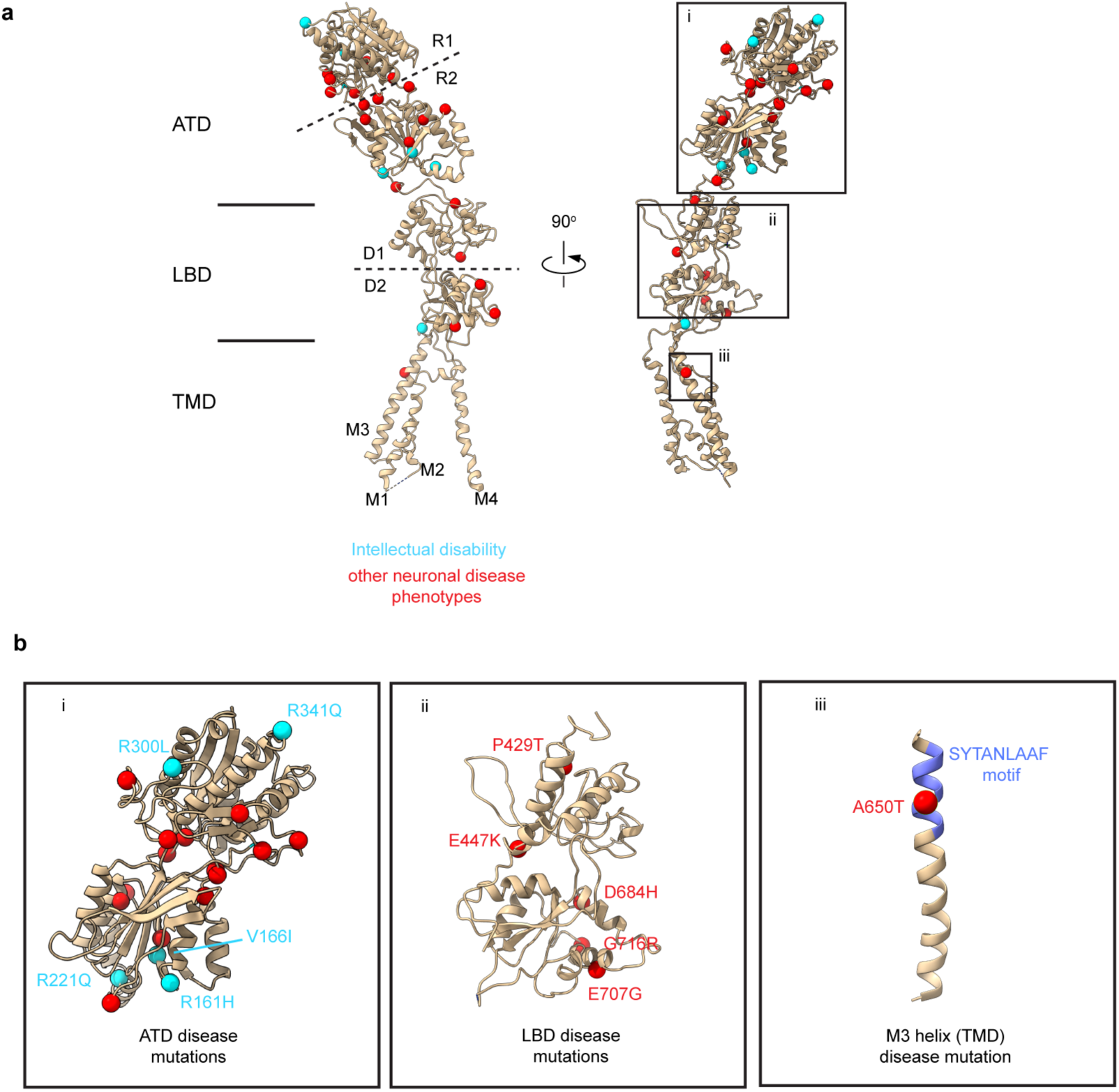
Disease-associated mutants and variants of hGluD1. **a.** Mapping of disease-associated mutants/variants onto hGluD1 structure, compiled from the reference 20. **b.** Insets showing highlighted domain-wise regions of interest (i) ATD, (ii) LBD and (iii) TMD.

**Extended Data Figure 8.**
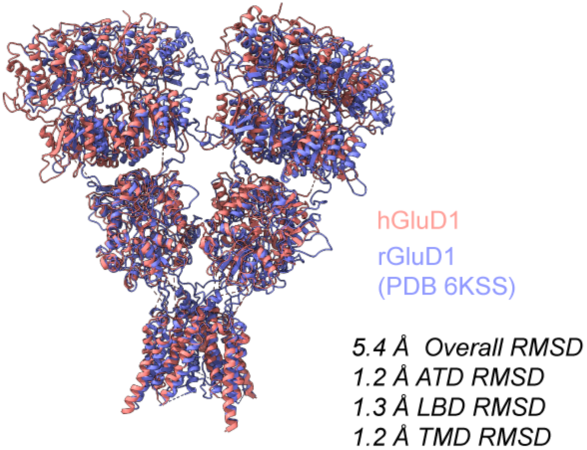
Structural Alignment. **a.** Structural alignment between hGluD1 (1mM Ca^2+^) and rGluD1 (PDB 6KSS).

**Extended Data Table 1:** Cryo-EM data collection, refinement and validation statistics.

|  | **hGluD1**<br>(EMD-xxxx)<br>(PDB xxxx) | **hGluD1<sub>Ca</sub>**<br>(EMD-xxxx)<br>(PDB xxxx) |
| --- | --- | --- |
| **Data collection and processing** |  |  |
| Magnification | 120,000x | 130,000x |
| Voltage (kV) | 200 | 300 |
| Electron exposure (e-/Å<sup>2</sup>) | 40 | 40 |
| Defocus range (μm) | -0.8 -2.5 | -0.8 -2.5 |
| Pixel size (Å) | 1.2 | .93 |
| Symmetry imposed | C1 | C2 |
| Initial particle images (no.) | 587,514 | 1,849,498 |
| Final particle images (no.) | 103,940 | 194,885 |
| Map resolution (Å) | 7.6 | 5.86 |
| FSC threshold | 0.143 | 0.143 |
| **Refinement** |  |  |
| Initial model used (PDB code) | hGluD1<sub>Ca</sub> | AF-Q9ULK0-F1 |
| Map sharpening *B* factor (Å<sup>2</sup>) | -484.7 | -462.5 |
| **Model composition** |  |  |
| Non-hydrogen atoms | 25704 | 25704 |
| Protein residues | 3268 | 3268 |
| Ligands | 0 | 0 |
| ***B* factors (Å<sup>2</sup>)** |  |  |
| Protein | 42.97/244.96/131.92 | 42.97/244.96/131.92 |
| Ligand | N/A | N/A |
| **R.m.s. deviations** |  |  |
| Bond lengths (Å) | 0.13 | .003 |
| Bond angles (°) | 0.924 | .586 |
| **Validation** |  |  |
| MolProbity score | 2.01 | 1.83 |
| Clashscore | 11.27 | 6.89 |
| Poor rotamers (%) | 0 | 0 |
| **Ramachandran plot** |  |  |
| Favored (%) | 93.06 | 92.99 |
| Allowed (%) | 6.78 | 6.92 |
| Disallowed (%) | 0.15 | 0.09 |

